# The complete genome and comparative analysis of a new *Tequintavirus*: *Salmonella phage Tennessee* Salten

**DOI:** 10.1101/2025.06.16.659957

**Authors:** Amandine Maurin, Carlos Zarate-Chaves, Cécile Breyton, Jacques Dainat, Alexandre Feugier, Rémy Froissart

## Abstract

The rise in multidrug-resistant pathogenic bacteria presents a major current challenge, highlighting the urgent need for alternatives and sustainable biocontrol strategies. Here, we report the genome analysis of a bacteriophage called *Salmonella phage Tennessee* Salten and attributed to a new *Tequintavirus* species. Salten was isolated following infection of *Salmonella enterica* subsp. *enterica* serotype Tennessee (sequence type ST5018). Its genome is 109,999 bp in length and contains 220 predicted proteins on which 197 are CDS and 23 tRNAs. Compared to its closest known relative phage *Escherichia phage* HildyBeyeler - sharing 84.6% identity - Salten harbours 16 unique or highly divergent genes. Of these, 13 encode proteins with unknown function, one encodes for a putative adenine methyltransferase and two encode HNH homing endonucleases. Moreover, the Long Tail Fibre protein, whose structure was predicted based on that of phage T5, was highly divergent among the *Tequintavirus* genus.

## Introduction

Nontyphoidal *Salmonella* infections represent a major foodborne public health concern, ranking as the second leading cause of bacterial outbreaks in humans and frequently associated with direct animal contact. *Salmonella enterica* is an *Enterobacteriaceae* that is responsible for a substantial burden of gastroenteritis in both humans and animals, with more than 93.8 million cases and approximately 155,000 deaths reported per year (Eng et al., 2015). The European Food Safety Authority (EFSA), the European Centre for Disease Prevention and Control (ECDC) and the U.S. Centers for Disease Control and Prevention (CDC) classified *S. enterica* as the second most reported-causing zoonoses (The European Union One Health 2023 Zoonoses report, 2024) and foodborne disease (Tack et al., 2019). *Salmonella* is a pathogen of global concern due to its ubiquity and persistence in diverse environments, including food processing facilities and agricultural settings. The extensive use of antibiotics as standard treatments in domains such as agriculture, livestock husbandry and food-processing further accelerates the emergence of resistant genotypes. The increasing prevalence of antimicrobial resistance (AMR) among pathogenic bacteria is a growing threat (Murray et al., 2022), compromising the efficacy of antibiotics or sanitary agents. *Salmonella enterica* is well known for its capacity to generate (multi)resistance genotypes, against both sanitary surface treatment and antibiotics (Curiao et al. 2016). As current sanitation and antimicrobial measures - particularly in the food industry - are becoming inadequate to fully eradicate *Salmonella* contaminations, it is an urgent necessity to consider alternative and sustainable methods for pathogen control.

One promising strategy for bacterial biocontrol is to reconsider the use of bacteriophages (phages), viruses that exclusively replicate within bacterial cells. This strategy favors the use of virulent phages over temperate ones, because (i) of their inherent ability to lyse bacterial hosts and (ii) to avoid the risk of transferring virulent factor or antibiotic resistance genes, which is a known risk with temperate phages (Kuhl et al., 2012; Balcazar 2014; Brown-Jaque et al., 2015).

We thus aimed to isolate a virulent phage for the biocontrol of *Salmonella enterica* subsp. *enterica* serotype Tennessee ST5018, a contaminating pathogen isolated from a pet food-industry. We reported here the analysis of the complete genome of *Salmonella phage Tennessee* Salten, a member of a new *Tequintavirus* species. We made several genome comparisons between Salten’s genome and (i) it’s closest relative *Escherichia phage* HildyBeyeler, (ii) four representative *Tequintavirus* genomes and (iii) more globally to 184 published genomes from the *Tequintavirus* genus. Notably, the Long Tail Fibre - implicated in the reversible adsorption on bacterial surfaces - was highly divergent among all the representative Tequintaviruses.

## Materials & Methods

### Phage isolation

The phage Salten was isolated from a bacterial strain of *S. enterica enterica* (https://enterobase.warwick.ac.uk/; accession number SAL-QB8962AA – named Sten2) serotyped as Tennessee by Eurofins (Aix en Provence, France). Sten2 came from a collection of samples obtained by swabbing a food-processing factory in Poland, over a two-year period (2017-2019). The French National Reference Center for *Escherichia coli*, *Shigella* and *Salmonella* at the Institut Pasteur (Paris, France) attributed the Sequence Type (ST) ST5018 to Sten2. Bacteria were routinely cultured in Lysogeny Broth (LB Lennox, Athena Enzyme Systems; Baltimore, MD, USA) or LB agar (1.2%).

A sample from Marseille’s wastewater (November 2017; 43°16′13″N, 5°24′00″E), filtered through a 0.22 µm Minisart polyethersulfone (PES) filter (#16541--K; Sartorius, Göttingen, Germany), and stored in glass bottles, was at the origin of the Salten isolate. Phage detection was done in a 96-deepwell plate, where each well contained 500 µL of LB inoculated with 2 µL of an overnight culture of Sten2 and 50 µL of filtered wastewater. The plate was incubated overnight at 37 °C within a ventilated incubator (AL 265-5; Aqua-lytic) with 450 rpm shaking (1.5 mm orbital; Titrama× 101 #544-11300-00; Heidolph Instruments, Schwabach, Germany). The following day, 50µL chloroform was added to each well and the plate was incubated at 4 °C for at least four hours. From the supernatant, 2 µL were transferred to a new 96-well polystyrene plate (#82.1581001; Sarstedt, Nümbrecht, Germany), containing 200 µL of LB supplemented in 10mM CaCl2 (Sigma-Aldrich, #C3881) and inoculated with 2 µL of an overnight culture of Sten2. Bacterial growth was monitored by evaluating turbidity through the measure of Optical Density at 600 nm wave length (OD600nm) over 16h-20h at 37°C with 300 rpm shaking (spectrophotometer FLUOstar Omega, BMG Labtech, Ortenberg, Germany).

The solution from the well showing the most delayed bacterial growth was transferred to a polypropylene 1,5 mL tube (Eppendorf SE, Hamburg, Germany). Residual bacteria were cleared by adding 10 % chloroform and centrifugation (10 min at 15,871 Relative Centrifugal Force or rcf, Eppendorf 5415 R) to keep only phages. The phage strain was then purified using the double-layer method (Kropinski et al., 2009). Briefly, 100 µL of the appropriate dilution of the phage solution was mixed with top LB agar (6g/L; #LF611001 Liofilchem, Italy) previously mixed with 100 µL of Sten2 overnight culture. After an overnight incubation at 35°C, one isolated lysis plaque was collected from the top agar, transferred into 200 µL SM buffer (100 mM NaCl, 10 mM MgSO_4_, 50 mM Tris-HCl, pH = 7.4) and incubated at 4 °C for at least one hour. The phage was then purified through five consecutive rounds of the double-layer method, picking one isolated lysis plaque at each round. The last round, the entire top LB agar layer was collected in SM buffer, centrifuged (10 min, 3,000 rcf, Eppendorf Centrifuge 5702R), filtered through 0.22 µm PES filter, and stored at 4 °C in polypropylen 15 mL tubes (#352096; Falcon, Corning, Mexico). The phage strain was named “*Salmonella phage Tennessee* Salten”, and hereafter called “Salten”.

### Transmission electron microscopy

Salten solution (15 mL at 10^11^ PFU/mL) was concentrated into 1.5 mL tubes by two rounds of centrifugation of one hour at 16,000 rcf, 4°C (Eppendorf Centrifuge 5415R). The pellet was resuspended in 600 µL ammonium acetate (100 mM; Sigma-Aldrich) and filtered through a 0.22 µm PES filter. Phages were then adsorbed onto a Formvar/carbon 300 grid (# CU 50/BX 9012.90.0000; Electron Microscopy Sciences, Hatfield, PA, USA), contrasted with 2% uranyl acetate, and visualized via transmission electronic microscopy (TEM, JEM-1400Plus, JEOL, Akishima, Tokyo, Japan).

### DNA extraction, preparation and sequencing

Phage DNA extraction was done according to a protocol (Gendre et al., 2022) adapted by Nicolas Ginet (Bacterial chemistry Laboratory, Marseille, France) after amplification of Salten in a more susceptible *S. enterica* Tennessee ST5018 isolate, *i.e.* Sten17 (Accession number SALQB8961AA in EnteroBase). Briefly, genetic material from bacterial origin potentially surrounding phages was eliminated by adding 10 µL of DNAse I (1 U/µL; #D5307; Sigma-Aldrich), 5 µL of RNAse A (10mg/mL; #EN0531; Thermo Fisher) and 2 µL of Dpn I (10 U/µL; #ER1702; Thermo Fisher). Phage DNA was extracted using phenol-chloroform-isoamyl acid 25/24/1 (#77617; Sigma-Aldrich). After DNA quantification with NanodropOne (Thermo Fischer) and Qubit 4 Fluoremeter (Invitrogen, Thermo Fisher), phage DNA was fragmented with a transposase enzyme. Fragmented end-prepared DNA was ligated to Illumina adaptors and then sorted with beads to select fragment sizes between 150-250 bp, for a final fragment size between 270-370 bp. Concentration of the final library was evaluated through Qubit 4 and fragment sizes were checked by migration using QIAxcel Advanced Instrument (QIAgen, Hilden, Germany). Fragments ready for sequencing sized between 280bp and 320bp. DNA libraries were sequenced in paired-ends in our in-lab sequencer (iSeq100; Illumina). Raw reads were deposited in the European Nucleotide Archive (https://www.ebi.ac.uk/ena/browser/support) with the following Accession Number: ERR13191102 (*Salmonella Phage Tennessee*).

### *De novo* phage assembly and annotation

After Illumina sequencing, phage reads quality was controlled using FastQC v.0.12.1 (Babraham Bioinformatics; https://www.bioinformatics.babraham.ac.uk/projects/fastqc/). Primers were trimmed with Fastp v.0.22.0 (Chen et al., 2018) using default parameters. Phage genome contigs were prepared following the workflow recommended in Turner et al., (2021). *De novo* phage genome assembly was carried out with SPAdes v.3.14.1 (Antipov et al., 2020; Prjibelski et al., 2020) with default parameters. We obtained a large contig of 110,076 nt with high coverage (average 400 reads depth), and a number of short contigs (below 1200 nt) with low coverage (average 10-20 reads depth). We kept only the largest Salten contig and generated a complete genome with PhageTerm Virome v.4.3 (Garneau et al., 2017). After polishing with Pilon v.1.24 (Walker et al., 2014), we obtained a new contig with a sequencing coverage ranged from 295× to 1171×. This contig corresponding to the complete genome was deposited in the European Nucleotide Archive (https://www.ebi.ac.uk/ena/browser/support) with Accession Number OZ075147.

Salten genome was annotated thanks to the Genome Annotation online tool from the Bacterial and Viral Bioinformatics Resource Center (Olson et al., 2023; BV-BRC, https://www.bv-brc.org/) using VIGOR4 v.4.0. and based on *Tequintavirus* annotation [Taxonomy ID = 187218]). Pharokka v.1.7.5 (Bouras et al., 2023) was used to complete the annotation. Structural genes were annotated manually based on Zivanovic et al (2014) and Linares et al (2023), using BLASTn or BLASTp from NCBI (Blast® services, available from: https://www.ncbi.nlm.nih.gov/Blast.cgi). The number attributed to each Coding DNA Sequences (CDS) was provided by the BV-BRC annotation. Proksee web server (Grant et al., 2023) was used to generate Salten linear genome map, through CGView builder v.2.0.5 (Stothard & Wishart, 2005). PhageScope web server 11 (https://phagescope.deepomics.org/) tools were used for lifestyle prediction, virulent factor and antimicrobial resistance gene detection.

### Comparative genomics analysis

The complete genome of the Salten was compared against the NCBI core nucleotide database using BLASTn. Hits with at least 30% query coverage were retained and, after genome comparison with VIRIDIC (Moraru et al., 2020), a subset of 184 phage genomes showing ≥70% average nucleotide identity (ANI) was selected for a refined analysis (*Tequintavirus* genus). Concerning the comparison of Salten to four other Tequintaviruses, gene coordinates were adjusted to standardize the starting point at the deoxynucleoside-5’-monophosphatase (dmp) gene for all genomes. VICTOR web platform was used to infer the phylogeny from nucleotides and using the D0 formula (Meier-Kolthoff & Göker, 2017). EasyFig (Sullivan et al., 2011) was used to generate detailed synteny diagrams. PanExplorer web server (Dereeper et al., 2022) was used to determine the core genome and Salten specific-genes (PanACoTA v.1.4.0 option with >80 % blast identity; Perrin & Rocha, 2021) by comparing to the 184 *Tequintavirus* genomes selected after VIRIDIC analysis.

A phylogenetic tree based on the aminoacid sequences of the Long Tail fibre proteins was performed based on a MUSCLE v.5.1 (Edgar, 2004) alignment followed by a tree construction through the Jukes-Cantor genetic distance model with neighbor-joining and 1000 bootstrap replicates, in Geneious Prime v.2025.1.3 (Kearse et al., 2012).

## Results and discussion

### Phage morphology

Transmission electron microscopy (TEM) of Salten revealed a T5-like Siphophage morphology, with a long non-contractile flexible tail of about 180 nm, attached to an icosahedral head of 60 nm of diameter (Fig. 1).

**Figure 1.**
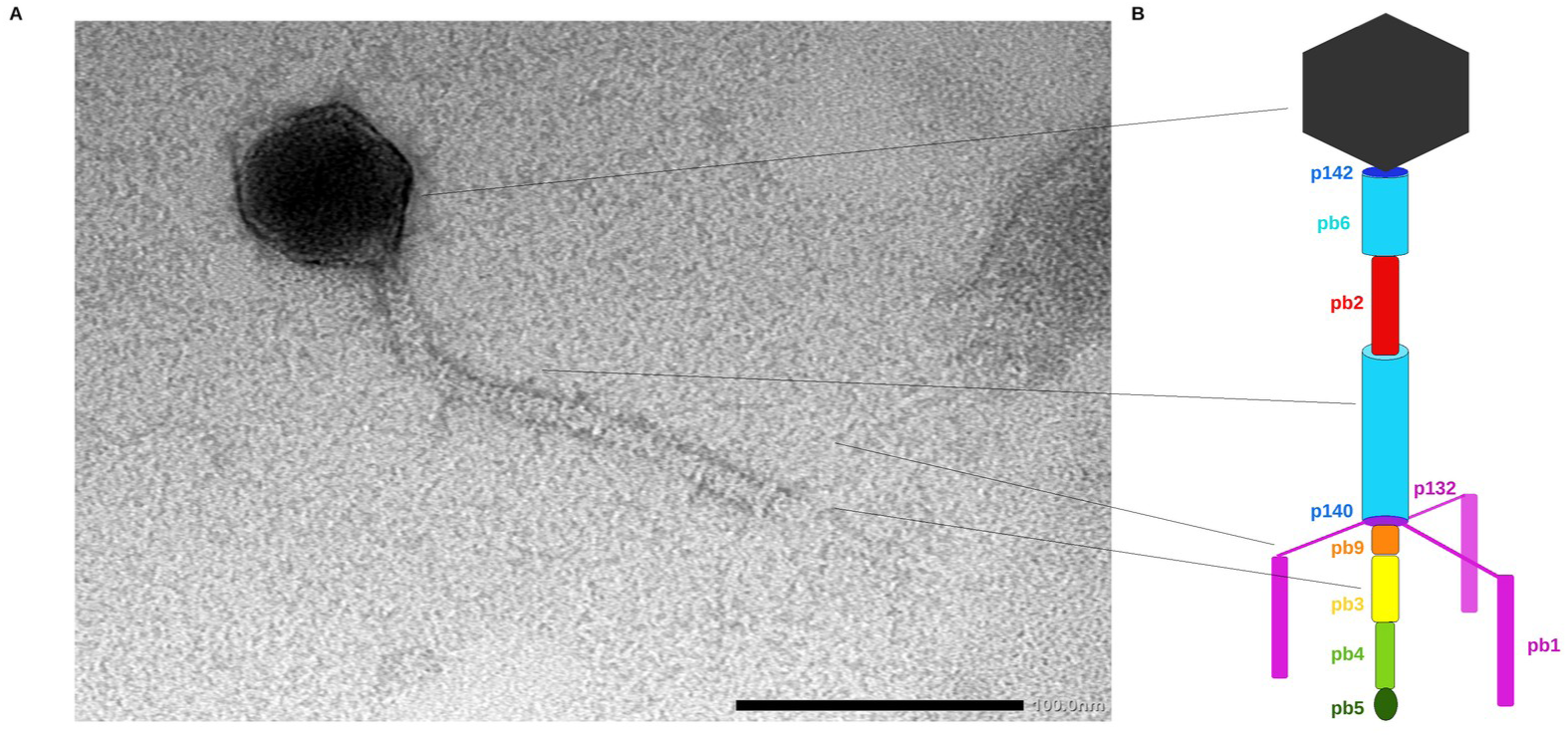
Morphology of the Tequintavirus Salten. **A-** Transmission electron microscopy (TEM). The scale bar represents 100 nm. **B-** Schematic structure of T5 adapted from Zivanovic et al. (2014) with TTPpb6: cyan; TMPpb2: red; p140 & p142: blue; pb9: orange; pb3: yellow; pb4: green; p132 & LTFpb1: pink; RBPpb5: dark green.

### Phage genome and features

Salten harboured a complete genome of 109,999 base pairs with a ∼39% GC content, varying across the genome (Fig. 2). A total of 220 Open Reading Frames (ORF) were predicted, including 197 CDS (Table S1) and 23 tRNAs. According to PhageScope, Salten was predicted to be virulent (lytic cycle) and did not encode any genes associated with virulence or antimicrobial resistance.

**Figure 2.**
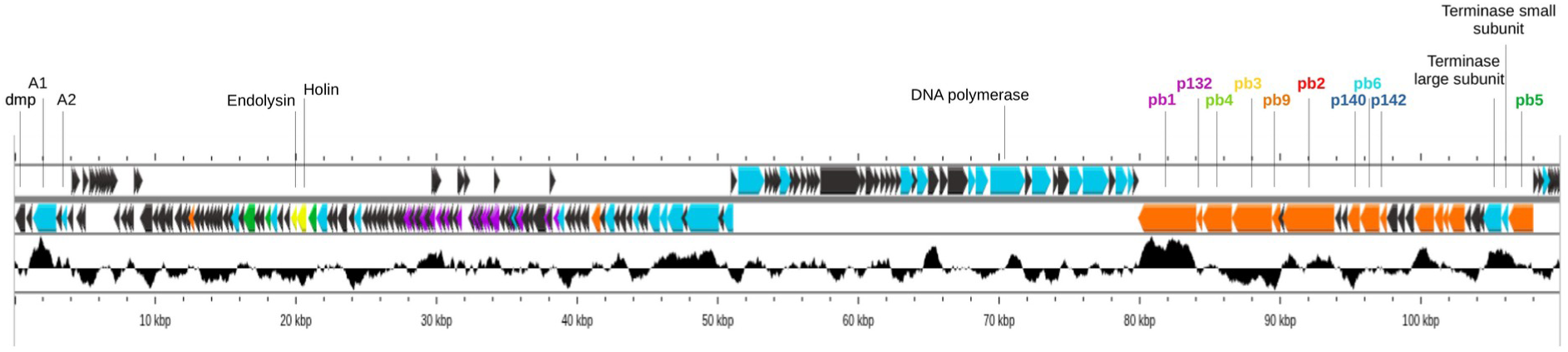
Genome linear map of Salten. GC content is indicated in black between the genome length and the predicted ORFs represented in arrows, where the direction indicates the forward (left to right) or reverse (right to left) strand of transcription. ORFs in light blue are associated to DNA transformation, in orange to structural proteins, in green to proteins modification, in yellow to bacterial cell-wall perforation and in purple to tRNA. Structural tail proteins names are colored according to the Figure 1.

The BV-BRC platform and pharokka annotated functional genes attributed to *Tequintavirus* and encoding for diverse proteins involved in host lysis, DNA replication, reparation and modification, protein modifications, *Tequintavirus* structure (tail and capsid proteins) and DNA packaging.

Salten contained the three best described pre-early genes, coding for the dmp, A1 and A2 proteins (CDS1, CDS3 and CDS5 respectively), which are essential for the second step of DNA insertion inside the host cell. The dmp and A1 are also involved in host DNA degradation, thanks to the phosphatase activity of the dmp which hydrolyse nucleotides in nucleosides-phosphate and the endonuclease activity of A1 (Davison 2015).

Salten encoded some proteins implicated in the perforation of the host membrane, such as a holin (CDS55) which creates pores inside the membrane and allow the entry of an endolysin (CDS54) for peptidoglycan cell-wall degradation from the inside of the host, in order to let out the new virions (Young 2013).

Salten contained several CDS encoding for proteins related to its own DNA transcription, replication and repair. Such autonomy for its own DNA replication facilitates a rapid replication and host cell lysis. We thus found CDSs encoding for enzymes involved in nucleotides metabolism (Wang et al., 2005), such as a dNMP kinase (CDS57), thymidylate synthase and dihydrofolate reductase participating to the transformation of dUMP in dTMP (CDS118, CDS119) and two ribonucleoside diphosphate reductases, one functioning in aeroby and the other in anaeroby (aeroby: CDS120 – small subunit, CDS122 – large subunit; anaeroby: CDS126). They reduce ribonucleoside diphosphates in desoxyribonucleoside diphosphates, which are the direct precursors of desoxyribonucleotides triphosphates (dNTPs; Hogenkamp 1983). Concerning DNA transcription and replication, Salten encoded for three helicases (CDS140, CDS151 and CDS156), two DNA ligases (CDS146 and CDS148), a primase (CDS153), an endonuclease (CDS162) and a DNA polymerase (CDS154). It contained a CDS encoding for the antitermination protein Q (CDS96), allowing the transcription of late genes (Grayhack & Roberts, 1982). We found a phage-associated recombinase (CDS159) and an exonuclease (CDS160) which share structural similarity with the Mre11 - Rad50 complex in eukaryotes or SbcCD in prokaryotes, gp46/gp47 in phage T4 and gpD13/gpD12 in phageT5. These complexes are involved in the detection and repair of double-stranded DNA breaks, contributing to genome stability and facilitating homologous recombination during viral replication (Rojowska 2013; Käshammer et al., 2019). CDS163 was annotated as a deoxyuridine 5’-triphosphate nucleotidohydrolase (dUTPase), a protein responsible for removing excess of dUTP mistakenly incorporated into DNA by DNA polymerases (Warner et al., 1979). Salten also encoded three transcriptional regulators (CDS141, CDS144 and CDS149) to control genes expression and two DNA-binding proteins (CDS142 and CDS158).

Salten encoded also two terminase proteins (large - CDS187 - and small - CDS188 - subunits) allowing DNA packaging (Casjens & Gilcrease 2009).

Other proteins are linked to proteins maturation or inactivation, such as a putative serine/threonine protein phosphatase (CDS47) and the ATP-dependent Clp protease (CDS56) which could degrade specific host or viral proteins (Shi et al., 1998; Kim et al., 2001). We found also a thioredoxin (CDS50) potentially implicated in enzyme maturation.

Between 79,875 bp and 107,981 bp, Salten encoded CDS producing structural proteins and DNA packaging. The capsid is encoded by two CDSs (CDS180 and CDS181) and the portal protein by CDS183. Thanks to the well modeled structure of T5 tail (Linares et al., 2023), we manually annotated the CDSs encoding for Salten’s tail proteins (Fig. 1B). The Tail Tube Protein pb6 (TTPpb6; CDS175) is surrounded by the Tape Measure Protein pb2 (TMPpb2; CDS171), which determines the length of the tube. Three lateral Long Tail fibres (each fiber being formed by the LTFpb1; CDS165) are connected by their amino-terminal (N-ter) part to the TTPpb6 protein through three lateral fibres p132 (CDS166). Anchoring of the LTFpb1 onto the central fibre is followed by the sequential positioning of pb9 (CDS169), pb4 (CDS167), pb3 (CDS168), and finally the receptor-binding protein pb5 (RBPpb5; CDS189) at the tip. RBPpb5 binds to outer membrane transporter, such as FhuA, FepA or BtuB, to trigger infection (Degroux et al., 2023). Small proteins p140 and p142 are encoded by CDS174 and CDS176, respectively.

### Comparative genomics with other Tequintaviruses

After complete genome comparison (VIRIDIC and BLASTn), *Escherichia phage* HildyBeyeler (MZ501074.1) was found to be the most similar to Salten, harbouring 84.6% average nucleotide identity (ANI). Based on the criterion that strains of the same species share at least 95% genome sequence identity (Adriaenssens and Brister, 2017), Salten can be considered as a member from a new species within the *Tequintavirus* genus. This statement was confirmed by the phylogenetic tree generated by VICTOR that classified HildyBeyeler and Salten in two different species (Fig. S1). Salten and HildyBeyeler shared 97.2% nucleotide identity within the core genome (i.e. CDS having >80% blast identity). Salten harboured 16 CDS (CDS7, 8, 9, 10, 11, 13, 28, 33, 51, 62, 67, 84, 85, 101, 116 and 127) absent in HildyBeyeler, of which 11 were annotated as “hypothetical proteins” and two as “Phage protein” (CDS33 and CDS127). CDS13 was annotated as a DNA N-6-adenine methyltransferase (*Salmonella phage* vB_SalS-SIY1lw [WVH10139.1]; 93.9% identity on 89% coverage), while the two last CDS (CDS51 and CDS62) were annotated as “Phage HNH homing endonuclease”.

VIRIDIC was used to delimit the *Tequintavirus* genus (i.e. >70% ANI) among NCBI phage accessions. To compare Salten to the diversity of Tequintaviruses, we selected four “representative” genomes out of the 184 genomes from the *Tequintavirus* genus. Selection was based on an interval of approximately 5% of ANI divergence, relative to Salten (since this threshold has been proposed as a species level of diversification; Adriaenssens and Brister, 2017). We thus selected *Escherichia phage* HildyBeyeler, *Salmonella phage* 8sent1748 (MT653146.1), *Escherichia phage* T5 (AY543070.1) and *Salmonella phage* S147 (NC_048012.1) with 84.6%, 80.4%, 76% and 70.1% ANI values, respectively. Genome comparisons performed with MAUVE showed a high degree of synteny (Fig. 3A; Darling et al., 2004), with three large collinear blocks, shared by all the genomes and kept in the same order. One of the HNH homing endonucleases (CDS62) - found in Salten and absent in HildyBeyeler - was also present in *Salmonella* phage S147, which was seen as a small collinear block in a different position of the genome (in dark green; Fig 3A). Knowing that HNH homing endonuclease are mobile genetic elements able to move by their own (Hafez and Hausner 2012, Stoddard 2014, Barth et al. 2023), it is not surprising to retrieve this CDS and its flanking sequences in different parts of the genomes.

**Figure 3.**
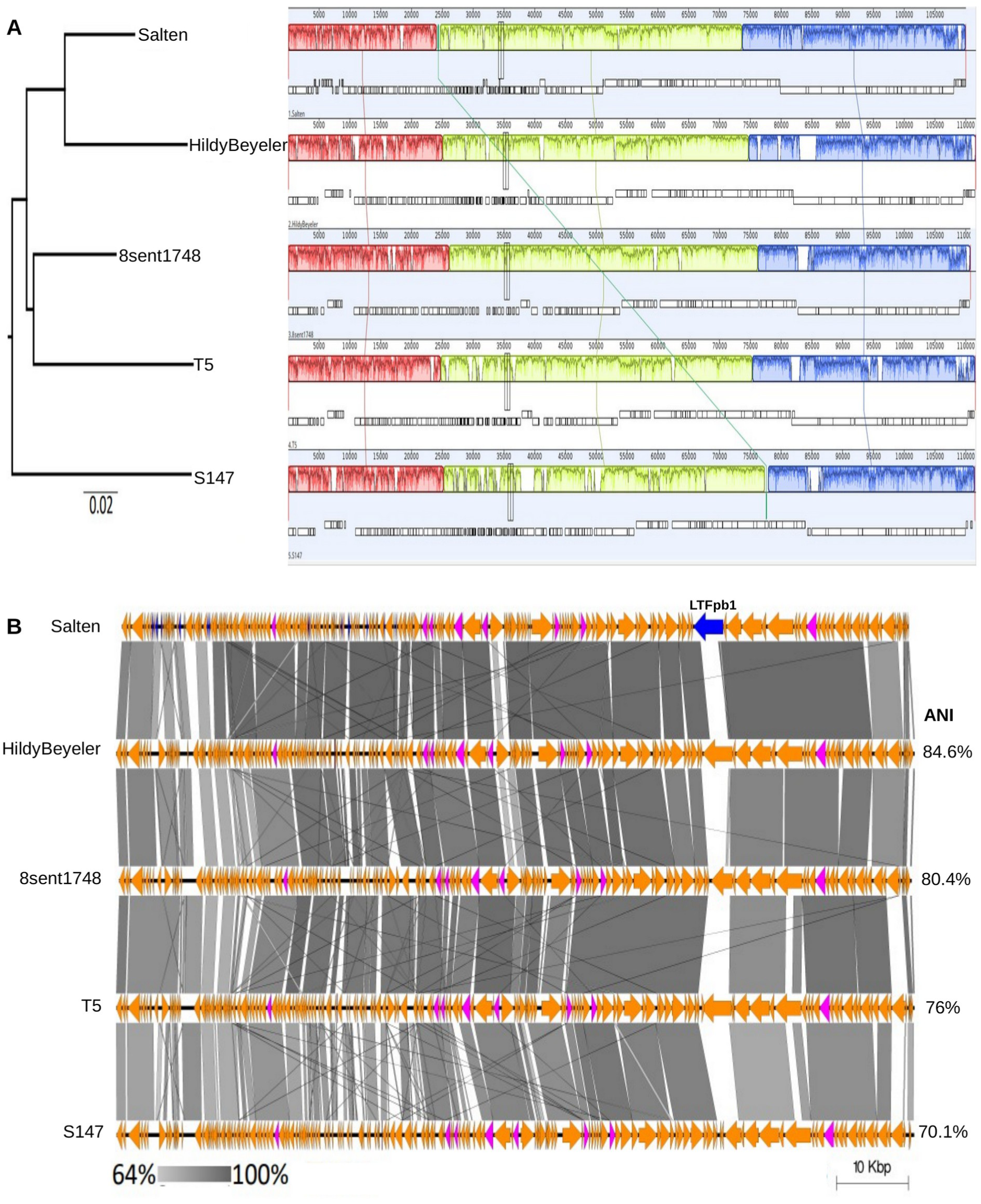
Comparative genomics analysis between Salten and the four representative Tequintaviruses. **A-** MAUVE alignment of five *Tequintavirus* genomes. The phylogenetic tree was built by VICTOR using the d0 formula and then re-arranged to have Salten on the upper part. **B-** EasyFig was used to obtain a more detailed synteny diagram. Lines connecting the genome maps indicate gene-level identity, ranging from 64% to 100%, represented by varying shades of grey. According to PanExplorer, genes in pink are homologs (>80% blastp identity) in the 185 *Tequintavirus* genomes, whereas genes in blue are unique to Salten or highly divergent. The percentage of ANI between each representative *Tequintavirus* genome and Salten is indicated in the right.

The pangenome of the five phages corresponded to 374 CDS (PanACoTA analysis). Among these pangenomic CDS, 64 of them were homologs between the five genomes - including Salten (>80% BLASTp identity), 121 CDS were shared by two or more genomes (and Salten harboured 80 of them) and 190 CDS were strain-specific (Salten harboured 54 of them). More globally, the pangenome of the 185 Tequintaviruses (including Salten) corresponded to 1472 CDS (PanACoTA analysis). Among these pangenomic CDS, only eight of them were homologs CDS (>80% BLASTp identity; pink on Fig. 3B; Table 1). These proteins can be considered essential for phage multiplication, such as DNA replication (dihydrofolate reductase and NAD-dependant DNA ligase) and protein degradation (metallopeptidase). The two genes annotated as “Phage proteins” (CDS113 and CDS140) have been compared to NCBI dataset (BLASTn) and CDS113 corresponded to a hypothetical protein or virion-structural protein (*Escherichia* phage phiLLS [YP_009790087.1]; 97.7% identity with 100% coverage) and CDS140 corresponded to a helicase (*Escherichia* phage DT57C [YP_009149867.1]; 99.6% identity with 100% coverage). When comparing Salten to all other 184 Tequintaviruses, 17 CDS were considered as unique to Salten (or highly divergent; represented in blue arrows in Fig. 3B) and were all annotated as “hypthetical proteins” or “Phage protein” (with no ability to improve annotation). To conclude, Salten as well as *Tequintavirus* genomes showed a high degree of variation and strain-specific CDS (accessory genome).

**Table 1.** List of the core genes (>80% blastp identity) among the 185 *Tequintavirus* genomes and named according to Salten BV-BRC annotation.

| CDS number | Protein name |
| --- | --- |
| 56 | ATP-dependent Clp protease proteolytic subunit |
| 111 | Metallopeptidase phage-associated |
| 113 | Phage protein / structural protein |
| 120 | Ribonucleotide reductase of class Ia (aerobic) |
| 124 | Phage phosphate starvation-inducible protein |
| 140 | Phage protein / helicase |
| 148 | DNA ligase phage-associated |
| 175 | Phage major Tail Tube Protein TTPpb6 |

In line with Skutel et al. (2023) findings that compared 15 LTFpb1 of *Tequintavirus*, we detected hypervariability in LTFpb1 between the 185 *Tequintavirus* genomes. When compared to the four representative genomes (Fig. S2), Salten’s LTFpb1 globally shared less than 64% of aminoacid identity (blue arrow annotated “LTFpb1” on Salten genome map in Fig. 3B). Indeed, LTFpb1 amino-acid sequences of T5, HildyBeyeler, 8sent1748 and S147 respectively shared 85.4%, 85%, 51.1% and 41.7% identity (coverage 78%, 48%, 31% and 31%) with the one of Salten. The N-terminal part of the protein (phage central tail binding domain; residues 1 to 41 on consensus sequence; Fig. S2) which is known to attach LTFpb1 to the phage central tail tube by the collar with p132 (Zivanovic et al., 2014) was well conserved. Additionally, the coiled-coil domain (residues 42 to 225; Fig. S2), responsible for projecting the lateral fibre away from the central tube, was also relatively well conserved. Notably, in Salten LTFpb1, the coiled-coil region contains insertions (repeats), leading to an extended coiled-coil. On the C-terminal part of LTFpb1, the chaperone encoding domain was also well conserved. However, the lateral fibre encoding domain (downstream the coiled-coil region; residues 226 to 874) showed a high level of variability (Fig. S2 and Fig. 4B-C). The LTFpb1 phylogenetic analysis was not congruent with the analysis made with the complete genomes of the phages. Indeed, Salten’s LTFpb1 was phylogeneticaly closer to T5 than the one of HildyBeyeler, and these three proteins were similarly distinct to the ones of 8sent1748 and S147 (Fig. 4A; Jukes-Cantor genetic distance model with neighbor-joining). This phylogenetic analysis was consistent with the structural analysis. Indeed, the structure of the lateral fibres of both T5 - crystallized and well modeled by Garcia-Doval et al. (2015) - and Salten were similar, but distinct from S147 (Fig. 4B). We therefore superimposed the predicted lateral fibres structure of Salten with the ones of S147 to visualise structure divergence between the two clusters (Fig. 4C). As proposed by sequence alignments (Fig. S2), their C-terminal domains did not overlapped. Given that this domain is known to reversibly bind to the bacterial polysaccharide moiety of lipopolysaccharides (LPS; Garcia-Doval et al., 2015; Heller et al., 1984), it is likely involved in host specificity and may play a critical role in determining the phage host range.

**Figure 4.**
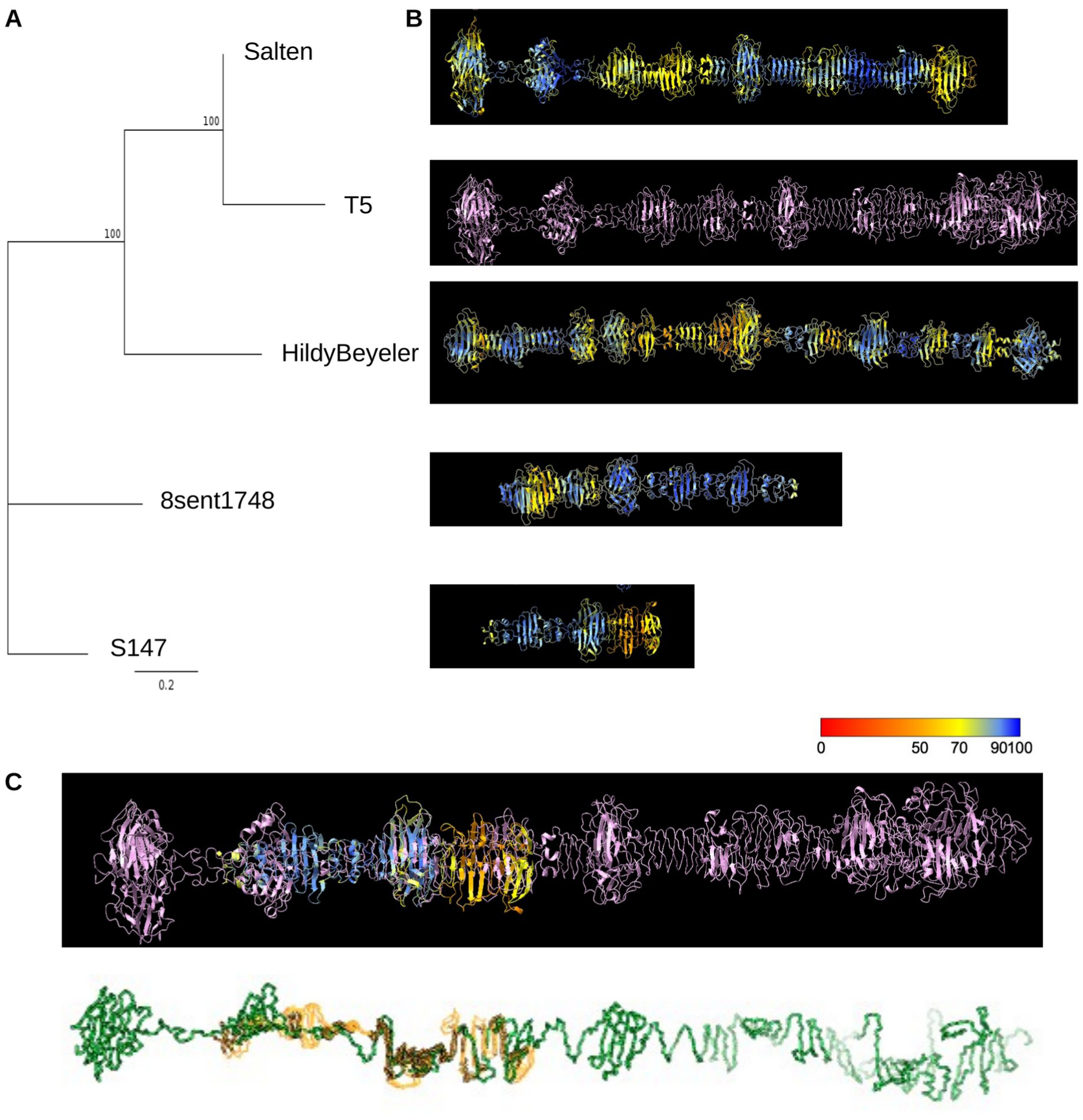
Comparative analysis of the LTFpb1 among the four representative *Tequintavirus* and Salten. **A-** LTFpb1 phylogenetic relationship reconstruction. LTFpb1 sequences were aligned with MUSCLE v3.8.425 in Geneious Prime 2025 and phylogenetic tree was constructed with Geneious Tree builder with Jukes-Cantor genetic distance model and neighbor-joining build method with bootstrap 1000. **B-** Structure of the predicted AlphaFold2 lateral fibre from Salten, T5 (pink), HildyBayeler, 8Sent and S147. Structures have been placed according to their position on the LTFpb1 phylogenetic tree. **C-** Top: superimposition of the lateral fibre of T5 LTFpb1 (pink) and S147 LTFpb1. Bottom: extended conformation of the lateral fibres structures to better visualise the overlay region between T5 (green) and S147 (yellow). Colors on the predicted structures from AlphaFold2 indicate the degree of confidence, from red (not very confident) to blue (really confident).

## Conclusion

The Salten bacteriophage was isolated from wastewater and exhibits a lytic activity against *Salmonella enterica* serotype Tennessee ST5018. It belongs to the *Tequintavirus* genus but remains the first member of a new species. As it does not harbour any bacterial virulence genes nor antimicrobial resistance genes, Salten is a promising candidate for the biocontrol of *Salmonella enterica* serotype Tennessee ST5018.

Notably, the LTFpb1 appeared to be highly diverse among the *Tequintavirus* genus. Given that this protein directly interacts with LPS on the host cell surface, such LTFpb1 diversity - particularly in the C-terminal domain - may explain Tequintaviruses host specificity. Indeed, O-antigen is known to be variable among Gram-negative bacteria (Goldman and Leive, 1980; Reeves and Cunneen, 2009). *Tequintavirus* adaptation to a new bacterial host may thus first reside in modification of their LTFpb1. Further studies would be interesting to determine which Salten’s aminoacids are involved in its interaction with its bacterial host. We could also experimentally investigate the capacity of Salten to increase its host range on other sequence types of *S. enterica* and related bacteria, as it was done for T3 phage and its gp17 fibre in Yehl and al. (2019).

## Acknowledgments

The authors thank M. Ansaldi (LCB, Marseille) for providing wastewater from which we isolated our phage, F-X. Weill for the bacterial sequencing, C. Mariac (DIADE, Montpellier) for the library quality check before sequencing and for QiAxcel use, A. Talman (MIVEGEC, Montpellier) for the help on the iSeq use, J. Garneau for PhageTerm help, M. Monot (I. Pasteur, Paris), A. Dereeper (PHIM, Montpellier), S. Bouzidi (MIVEGEC, Montpellier) and J. Hayer (MIVEGEC, Montpellier) for bioinformatics advices. We thank UMR MIVEGEC for providing an efficient working environment and acknowledge the ISO 9001 certified IRD itrop HPC (member of the South Green Platform) and the whole bioinformatics team support in Montpellier for providing HPC resources that have contributed to the research results reported within this paper. URL: https://bioinfo.ird.fr/ - http://www.southgreen.fr.

## Data availability

The data that support the findings of this study are openly available at https://src.koda.cnrs.fr/MAURINAmandine/salten_report

## Author contribution

Conceptualisation, R.F. and A.M.; Methodology & Investigation, R.F. and A.M. with the help of C.Z.-C. for the genomic comparison; Results analysis, R.F., A.M., C.Z.-C., C.B., and J.D.; Writing, R.F. and A.M.; Revision, R.F., A.M., C.Z.-C., C.B., and J.D..

